# ProteoCast: a web server to predict, validate, and interpret missense variant effects

**DOI:** 10.1101/2025.11.03.686214

**Authors:** Marina Abakarova, Maria Inés Freiberger, Arnaud Liehrmann, Michael Rera, Elodie Laine

**Affiliations:** Sorbonne Université, CNRS, IBPS, Department of Computational, Quantitative and Synthetic Biology (CQSB, UMR7238), 75005 Paris, France; Université Paris Cité, INSERM UMR U1284, 75004 Paris, France; Institut Universitaire de France (IUF).

## Abstract

Understanding how mutations affect protein function remains critical yet challenging, particularly for variants in clinical databases lacking experimental characterisation and for intrinsically disordered regions. Current computational approaches often operate as black boxes, providing predictions without sufficient transparency or quality assessment of the underlying data. Here we present ProteoCast, a user-friendly web server that predicts variant effects through evolutionary constraint analysis and structural context integration. ProteoCast provides a three-tier variant classification (impactful, mild, neutral) to help prioritise mutations for clinical interpretation and experimental validation. It incorporates multiple sequence alignment quality controls to ensure prediction reliability and flag positions with insufficient evolutionary information. Beyond single-variant classification, ProteoCast employs a novel segmentation approach based on mutational sensitivity to identify functional linear peptides in disordered regions. Interactive visualisations guide users through results interpretation, from variant-level predictions to protein-wide functional landscapes. Evaluation on 63,000 ClinVar variants demonstrates 77% sensitivity and 87% specificity for pathogenicity prediction, with performance maintained across species (85% accuracy on Drosophila lethal mutations). ProteoCast successfully identifies twice as many functional motifs in intrinsically disordered regions compared to conservation-based phylogenetic methods. Predictions can be tuned to specific conformations, such as bound forms in protein complexes, for improved accuracy and interpretability. With its transparent, unsupervised methodology and computational efficiency (minutes per protein), ProteoCast democratises access to variant effect prediction and functional site discovery for the broader research community. The web server is freely available at: https://proteocast.ijm.fr/.

**Highlights:**

- ProteoCast web server democratises variant effect prediction through a fast, transparent platform combining evolutionary analysis and structural context
- Three-tier variant classification enables prioritisation for clinical interpretation and genome editing experiments
- Novel mutational sensitivity segmentation approach identifies functional peptides in disordered regions
- Interactive visualisations and input alignment quality controls guide interpretation

## Introduction

Predicting the impact of point mutations on protein functions is a central challenge in protein engineering, clinical genomics, and genome editing. Systematically assessing these effects enables researchers to guide protein design, determine the clinical significance of patient variants, and prioritise targets for CRISPR experiments. In recent years, the massive expansion of publicly available protein-related data has driven a surge in interest in developing computational methods to address this challenge. Progress in the field can be monitored through the comprehensive benchmark ProteinGym, comprising over 2.5 million variants from 217 deep mutational assays^1^. While the number of Variant Effect Predictors (VEPs) has doubled since the first release in December 2023, the improvement has remained limited and unequal across the measured phenotypes, with notable gains for thermodynamic stability but negligible progress for organismal fitness prediction. Understanding these limitations requires examining the diversity of approaches.

VEPs vary widely in terms of input data – encompassing protein sequences, population variants, and three-dimensional (3D) structures, learning paradigm, complexity, and computational cost. Notably, relatively simple unsupervised heuristics relying on a few biologically meaningful hypotheses^2–4^ perform on-par with high-capacity deep learning architectures, namely protein Language Models (pLMs), trained on massive amounts of protein data. Scaling pLM beyond 1 billion parameters does not improve accuracy – performance even degrades beyond 4 billion parameters^5^. Moreover, pLMs taking single sequences as input underperform compared to those explicitely leveraging evolutionary information from multiple sequence alignments (MSAs) during training^6^ or at inference^7^, and to multimodal models incorporating structural knowledge^8,9^.

Desired properties for a VEP include computational efficiency, transparency of the algorithm and interpretability of the output, as well as the ability to capture subtle functional variant impacts. In this respect, intrinsically disordered regions (IDRs), whose functional roles are increasingly recognised^10,11^, are especially challenging^12,13^ due to two key factors: traditional structural features provide little insight on their function and mutational sensitivity, and IDRs exhibit high sequence divergence with weaker evolutionary constraints than folded domains^14,15^. Seminal works for identifying functional motifs within IDRs focused on the detection of “islands of conservation” in these rapidly evolving regions^16–18^. They have addressed this problem with relative local conservation metrics^18^ or two-state phylogenetic Hidden Markov Models^17^ aiming at contrasting individual columns in MSAs against a background context of 20-30 residues. Their detection rate remains relatively low, suggesting that many of these functional motifs lack strong conservation signals. Moreover, detection is highly sensitive to alignment quality, typically requiring carefully curated sets of a few dozen orthologous sequences. While pLMs are independent of MSAs and could in principle help decrypt IDR sequences, they currently offer no enhanced predictive power in IDRs^12^.

Here we present the ProteoCast web server, a user-friendly platform for predicting point mutation effects on protein function. ProteoCast builds upon our previously developed approach for computing evolutionary scores from MSAs^4^. It provides users with comprehensive quality controls of the input MSA and a three-tier classification of variants (impactful, mild, or neutral), enabling the distinction between mutation-sensitive and mutation-tolerant residues. Beyond variant classification, ProteoCast can optionally leverage protein structural data^19,20^ to refine predictions for specific conformations and enable structural mapping for mechanistic interpretation. In addition, it exploits mutational sensitivity—rather than simple conservation—combined with powerful segmentation approaches^21,22^ to identify putative functional linear peptides within IDRs.

ProteoCast is fully unsupervised, with a transparent methodology, and interpretable features. It runs in minutes for most proteins and provides interactive visualisations to help users gain insights into genotype-phenotype relationships. ProteoCast webserver is available at https://proteocast.ijm.fr/.

ProteoCast addresses the following key questions:

- Given a variant, is there sufficient evolutionary information about its position and protein context? To what extent is it counter-selected during evolution, and thus likely to disrupt protein function and impact organismal fitness?
- Given an unstructured protein region, which peptides or residues are most evolutionarily constrained and thus likely play essential functional roles? Which variants are tolerated at these positions?
- Given a 3D structure, which residues are under selective pressure while not being buried in the protein core, making them good candidates for modulating protein interactions and regulation? Given a protein complex, which variants are likely to directly alter binding?

## Materials and methods

### Capturing variant effects from evolutionary information

To predict variant effects, we integrated our previously developed Global Epistatic Model (GEMME)^4^ in the web server. GEMME computes evolutionary scores based on a measure of conservation accounting for the topology of the tree relating the sequences in the input MSA^23,24^ and on a minimal evolutionary distance that quantifies the amount of changes required to accommodate each mutation^4^ (**Fig. 1**, *Mutational Landscape*). The generated mutational landscape is screened for positions with low score dispersion. If the position is poorly conserved, and the input MSA is highly gapped or samples a small fraction of the 19 possible substitutions, then ProteoCast flags the position as non-confident (**Fig. 1**, *Residue Confidence*). ProteoCast then models the predicted evolutionary score distribution with a mixture of three Gaussians. The parameters of these Gaussians are used to define thresholds for classifying variants as impactful, mild, or neutral (**Fig. 1**, *Variant Classification*). In turn, residues where more than half of the possible substitutions are neutral are considered as tolerant to mutations, and the others sensitive (**Fig. 1**, *Residue Classification*). For more methodological details, see^25^.

**Figure 1.**
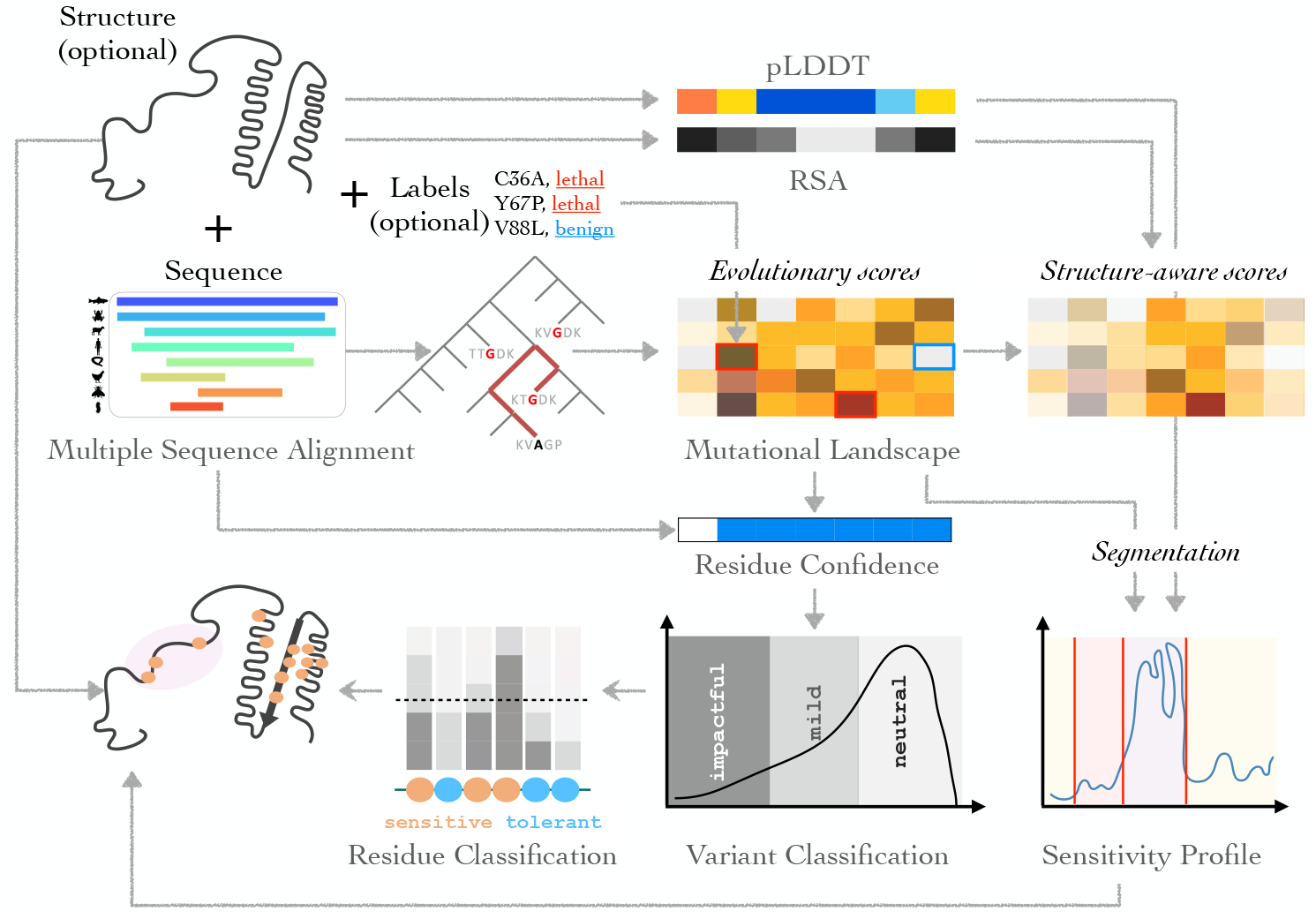
Overview of ProteoCast’s approach. ProteoCast takes as input a MSA with the ungapped query protein sequence on top, and, optionally, a 3D structure and/or a list of annotated mutants. It computes evolutionary scores for every possible substitution at every position in the query protein (*Mutational landscape*). It highlights the known mutations provided by the user on the landscape (one colour for each unique label). It flags potentially unreliable predictions based on a quality assessment of the input alignment (*Residue Confidence*). It automatically classifies variants into neutral, mild, or impactful based on the full raw score distribution (*Variant Classification*). It then determines whether each position in the query protein is sensitive or tolerant to mutations based on the proportion of mild and impactful substitutions (*Residue Classification*). In parallel, ProteoCast identifies linear protein segments with elevated sensitivity compared to their background, optionally using pLDDT-derived weights (*Sensitivity Profile*). In addition, it tunes the evolutionary scores to the input 3D conformation using its per-residue relative solvent accessibility (*RSA*). Finally, it maps the per-residue predictions onto the input 3D structure.

### Segmenting mutational sensitivity profiles

To identify putative regulatory and interacting peptides in unstructured regions, we detect changepoints in the protein’s mutational sensitivity profile (**Fig. 1**, *Sensitivity Profile*). The mutational sensitivity of a given residue is estimated as the average of the evolutionary scores computed for all 19 substitutions at that position. We model this measure as a piecewise-constant signal, where each segment contains observations drawn from independent Gaussian distributions sharing a common variance *σ*^2^ but with distinct segment-specific means *µ*_*j*_. This approach aims at identifying contiguous functional regions evolving under distinct selective regimes, in contrast with gamma-distributed rate models that assume continuous, site-independent variation. Such segment-based models have established precedent in molecular evolution and reflect the biological reality that proteins are modular entities with functionally distinct regions, including linear peptides embedded in intrinsically disordered regions^16–18^.

Parameter estimation relies on the FPOP dynamic programming algorithm^26,27^, which computes the penalised maximum likelihood through functional pruning—tracking the likelihood as a piecewise quadratic function of the final segment’s mean parameter and efficiently updating it as new observations are incorporated. The penalty term *ασ*^2^ log(*n*) controls segment number, with *α* = 1.4. Variance *σ*^2^ was estimated using the Hall estimator^28^. Since high-confidence AlphaFold2^29^ regions (predicted Local Distance Difference Test^30^ pLDDT > 70) exhibited greater mutational sensitivity variance that could lead to over-segmentation, we applied differential weighting (*w* = 0.1 for high-confidence regions, *w* = 1 elsewhere) to avoid spurious changepoint detection. Each resulting segment was assigned a discrete score (0, 1, or 2) characterising whether its mean falls below, between, or above the means of its flanking segments. We note that our approach is multiscale: it can detect segments ranging from single critical residues to entire domains, with optimal boundaries and lengths determined automatically from the evolutionary signal. The granularity of partitioning is tunable to the user’s specific needs.

### Sequence-Structure Alignment

To map residue positions between the input FASTA sequence and PDB structure, ProteoCast performs a global pairwise alignment using the Biopython *PairwiseAligner* module^31^. It first extracts the amino acid sequence of the chain of interest from the PDB file using the *gemmi* library^32^, retaining only standard residues. Then, the alignment is performed with the following scoring parameters: match score = 1.0, mismatch score = −3.0, gap opening penalty = −2.5, and terminal gap and gap extension penalty = −2.0.

### Modulating predictions with solvent accessibility

Relative Solvent Accessibility (RSA) quantifies the extent to which each amino acid residue is exposed to the solvent, expressed as the ratio of accessible surface area to the maximum possible surface area for that residue type (**Fig. 1**, *RSA*). ProteoCast computes per-residue RSA using the Shrake–Rupley algorithm^33^, as implemented in the Biopython module *Bio*.*PDB*.*SASA*.*ShrakeRupley* ^31^. Default parameters are used, including a probe radius of 1.4 ^°^A (representing a water molecule) and 100 sampling points per atom sphere. Non-standard amino acids (*e*.*g*., selenomethionine, methylated residues) are mapped to their standard equivalents via a curated dictionary (*AminoAcid. NON STANDARD AAS*) or ignored if there is no equivalent. This procedure was adapted from Tsishyn et *al*.^2^. For multimeric structures, ProteoCast computes the RSA of the chain of interest in the original context (complexed) and in isolation (free).

ProteoCast computes structure-aware variant scores for residues aligned between the input sequence and structure as,

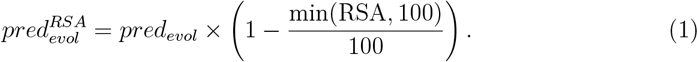

This formulation straightforwardly weights the evolutionary score *pred*_*evol*_ of a variant by the solvent accessibility of the wild-type residue, as proposed in ^2^ (**Fig. 1**, *Structure-aware scores*). The more exposed a residue, the smaller the weight. RSA values are capped at 100% to handle edge cases where calculated accessibility exceeds theoretical maximum values. Residues that cannot be structurally aligned are assigned NaN values and are explicitly displayed in grey and labeled as “unaligned” in the mutational landscape visualisations.

### Evaluating predictions

We evaluated the accuracy of ProteoCast variant classification against (1) ∼200 assays from the ProteinGym benchmark^1^, (2) ∼ 63,000 pathogenic and benign human variants from ClinVar^34^, (3) ∼1,000 mutations developmentally lethal in *D. melanogaster* from FlyBase^35^ and hundreds of thousands of fly population variants from DGRP^36^ and DEST2^37^, and (4) ∼500 yeast proteins with annotated functional sites from^17^. For (1), we retrieved the input MSAs and AlphaFold 3D from the ProteinGym benchmark^1^. For (2), we directly took the MSAs and evolutionary scores we computed in a previous study^38^ (https://doi.org/10.5061/dryad.vdncjsz1s) and applied the subsequent ProteoCast analyses to them. For comparison, we downloaded EVE bulk predictions from: https://evemodel.org/download/bulk. For (3), we generated Proteo-Cast predictions over the whole fly proteome from MSAs computed using ColabFold^39^ and AlphaFold 3D models either retrieved from AFDB or computed with ColabFold. For (4), we generated the input MSAs with the ColabFold protocol (uniref30 2302, colabfold envdb 202108) and used 3D models from AFDB. For comparison, we retrieved phylo-HMM predictions from the Supplementary Information of^17^. More details about (2-4) evaluations can be found in^25^.

### Web server implementation

ProteoCast webserver is a *Django*-based project running on a virtual machine hosted by *Institut Jacques Monod* (Dell PowerEdge R740 server). The backend was developed in Python using *Django*, while the frontend was built with HTML, CSS, and JavaScript to provide a user-friendly interface. The application was deployed on an Apache2 web server with Django’s WSGI integration. It incorporates *plotly* for interactive data visualisation and *Mol* (Molstar)*for 3D structure visualisation. Data handling and processing were facilitated using *pandas*. ProteoCast source code can be accessed at https://github.com/abakarovaMarina/ProteoCast.

### Web server description

ProteoCast predicts the impact of all possible single-point mutations in a query protein, identifies putative regulatory or binding sites in its unstructured regions, and maps the predictions on its 3D structure. The web server is freely available at https://proteocast.ijm.fr/.

### Submission page

The simplest way to use ProteoCast is to provide a UniProt accession code as input. The corresponding MSA and protein 3D structure will be automatically retrieved from the AlphaFold database (AFDB)^19^ (**Fig. 1**). Alternatively, users may provide a custom MSA in A2M, A3M, or FASTA format and/or a custom 3D structure in PDB format. The 3D structure is optional and is not required to match the query (first) sequence in the input MSA exactly. Partial coverage, alternative isoforms, or complexes are all supported. For multimeric structures, users specify the chain of interest (default: chain A). In addition, users may upload a list of annotated mutations in CSV format, specifying the wild-type amino acid, its position, the substituting amino acid, and a label, *e*.*g*., a known phenotype (pathogenic, benign, lethal, population,…etc).

### Result page

#### Comprehensive mutational scanning

ProteoCast generates predictions for all possible single amino acid substitutions across the protein sequence, yielding a complete mutational landscape of 19 × *L* variants, where *L* is the protein length (**Fig. 1**). It reports a *raw evolutionary score* for each variant, where a more negative score indicates stronger predicted impact, and a binary confidence value for each residue, where a null value flags position with insufficient evolutionary information in the input MSA. To further help user assess the input sequence diversity, ProteoCast shows a plot summarising the MSA where each line represents a sequence and is coloured according to its similarity to the query sequence. To facilitate interpretation, ProteoCast further classifies variants into neutral, mild, and impactful by fitting a three-component Gaussian mixture model (GMM) to the distribution of raw scores (**Fig. 1**, *Raw score distribution* panel). In addition, it computes *structure-aware scores* that incorporate relative solvent accessibility (*RSA*) from the provided 3D structure. The predicted raw evolutionary scores, classes, and structure-aware scores are displayed as interactive heatmaps.

#### Sensitivity Profile segmentation

The interactive mutational sensitivity profile displays the average evolutionary scores per residue along the protein sequence (**Fig. 1**). ProteoCast partitions the protein sequence into a set of segments exhibiting homogeneous mutational sensitivity through changepoint detection. The identified segments are coloured depending on whether they have elevated sensitivity compared to both neighbouring segments (purple) or only one (red). The granularity of the partitioning is locally modulated by AlphaFold pLDDT values extracted from the input 3D structure and displayed on top of the mutational sensitivity profile, enabling direct visual comparison between structural confidence and evolutionary constraint.

#### 3D visualization

The 3D structure visualisation displays the protein structure with interactive colouring schemes to represent different structural and evolutionary per-residue features (**Fig. 1**). When a UniProt identifier or PDB structure is provided by the user, the structure is rendered along with its experimental b-factors or pLDDT scores for AlphaFold models. Three alternative colouring schemes are proposed:

- **Sensitivity mode**: residues are coloured according to their mutational sensitivity.
- **Residue Class mode**: residues are coloured according to their class, either sensitive to mutations (more than half of the 19 substitutions are non-neutral) in red or tolerant in blue.
- **Structure-aware Sensitivity mode**: Residues are coloured according to structure-aware scores emphasising buried, constrained positions while down-weighting surface-exposed residues.

In all three modes, only the residues aligned with the query sequence are coloured, while the others are displayed in gray. An additional feature allows highlighting segments with elevated sensitivity located in unstructured regions (pLDDT < 70).

### Past Jobs page

Users can access results for any completed job by providing its identifier, *e*.*g*. 20251008122309, through the Past Job page.

## Results

### Classifying variants

ProteoCast demonstrates state-of-the-art performance in discriminating pathogenic from benign mutations in humans. Specifically, it achieves a per-protein average Area Under the Curve (AUC) of 0.923 against ∼ 63,000 variants annotated in ClinVar^34^, compared to 0.920 for PoET^40^ and TranceptEVE^41^, 0.919 for GEMME^4^, and 0.914 for EVE^42^, the top performing predictors in ProteinGym leaderboard. In addition to continuous scores, ProteoCast classifies variants as neutral, mild or impactful, based on their evolutionary constraints relative to the whole protein context (**Fig. 2A**). For this categorical classification task, ProteoCast exhibits similar performance to the well-established EVE method, with 81.9% sensitivity and 88.4% specificity compared to 86.6% and 88.6% for EVE on a subset of 33,274 variants for which EVE classification is available (comparison detailed in^25^). EVE classification does not cover the full ClinVar set because it applies stringent thresholds on MSA coverage and requires high probabilities from its fitted Gaussian Mixture Model. By contrast, ProteoCast can leverage sparser evolutionary information: it determined that 99% of all ClinVar variants have sufficient data for prediction and maintained 77.1% sensitivity and 86.9% specificity on this comprehensive set.

**Figure 2.**
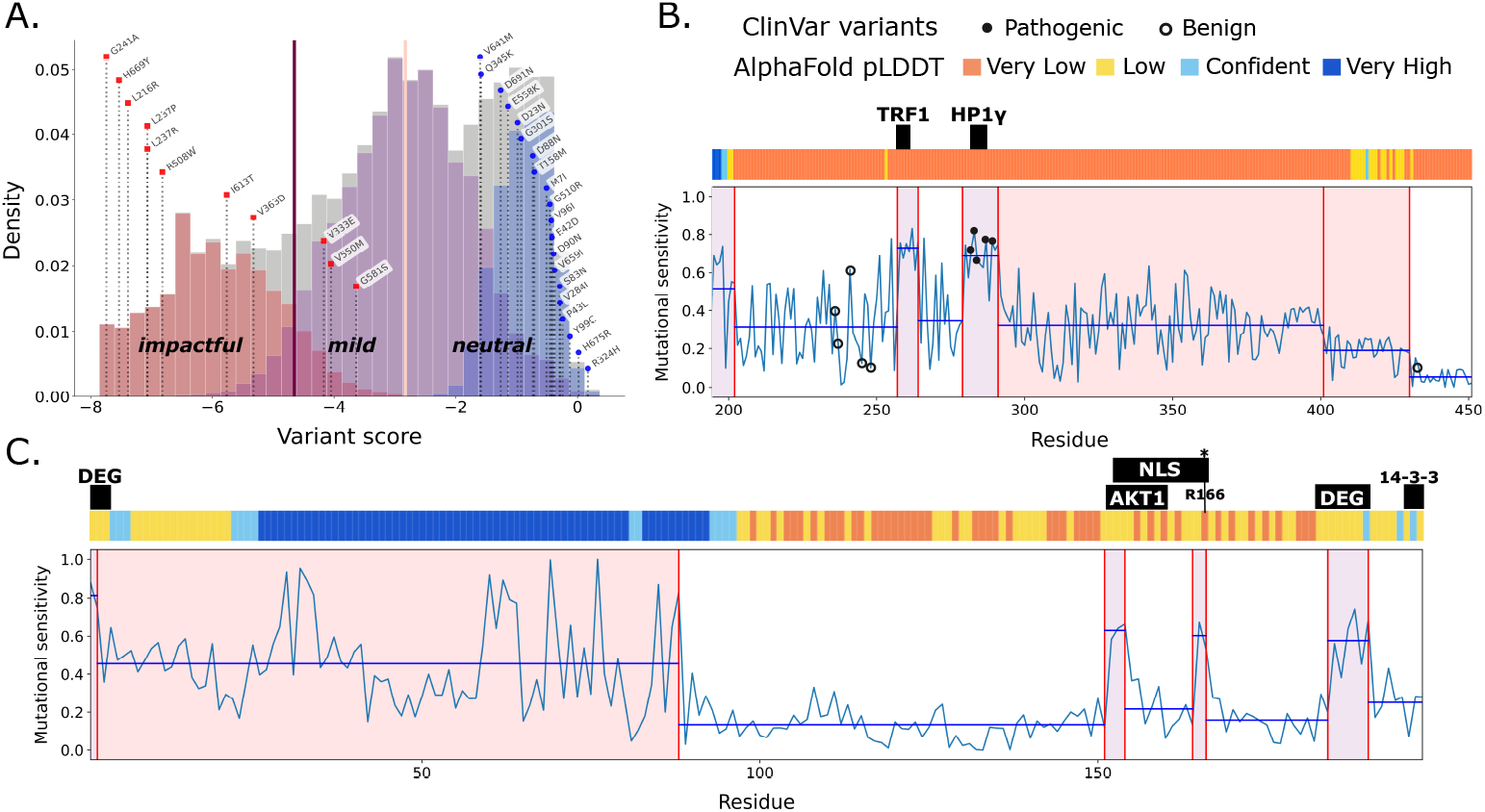
Variant classification and mutational sensitivity profile segmentation. **A**. Illustration of Proteocast’s variant classification using Biotin–protein ligase (UniProt id: P50747, ProteoCast job id: 20251008122309). Variants annotated in ClinVar as pathogenic (red) or benign (blue) are mapped on the predicted variant score distribution. The three fitted Gaussians are coloured in red, purple, and blue. The classification thresholds are depicted as vertical straight lines in purple and pink. **B-C**. Segmented mutational sensitivity profiles for the TERF1-interacting nuclear factor 2 TINF2 (B, UniProt id: Q9BSI4, ProteoCast job id: 20251009204919) and the cyclin-dependent kinase inhibitor p27^*Kip*1^ (C, UniProt id: P46527, ProteoCast job id: 20251010165911). The horizontal black lines indicate the mean sensitivity for each segment and segments with elevated sensitivity compared to their neighbours are coloured in purple (higher than both neighbours) or red (higher than one neighbour). The coloured bar on top gives the AlphaFold2 pLDDT. The black boxes indicate the functional linear peptides annotated in the Eukaryotic Linear Motif (ELM) database^43^ or known from the literature^44,45^. See also the results for KCNK3 (UniProt id: O14649, ProteoCast job id: 20251011164558) whose 14-3-3 binding motif is detected.

Beyond human genomics, ProteoCast generalises to other organisms such as *Drosophila*, where it detected 85% of ∼1,000 lethal mutations referenced in FlyBase as non-neutral, and it correctly identified 88% of ∼137,000 variants from inbred laboratory lines^36^ and 82% of ∼270,000 natural population variants^37^ as neutral. Together, these results demonstrate that ProteoCast enables variant prioritisation across species, from interpreting uncertain clinical variants in humans to guiding CRISPR experiments in model organisms. The three-tier classification provides additional granularity: impactful variants, under strongest evolutionary constraints, typically affect essential structural or catalytic residues, while mild variants likely have more subtle effects on protein stability, regulation and interactions. ProteoCast achieves these results without prior training on labelled variants and without variant filtering, making it broadly applicable across proteins and organisms.

Figure 2A illustrates ProteoCast’s variant classification on the Biotin–protein ligase: all 11 pathogenic mutations in ClinVar are assigned to the impactful or mild categories while all 20 benign mutations are correctly identified as neutral. This example was selected to clearly show the three-tier classification system: more complex cases where pathogenic and benign mutations overlap in their evolutionary constraints require more nuanced interpretation and are systematically evaluated in^25^. Notably, variants of unknown significance vastly outnumber classified variants for this protein (159 versus 31), illustrating the knowledge gap in variant interpretation. ProteoCast helps address this gap by classifying two thirds (105 out of 159) of these variants as neutral, 15% (23) as impactful and 19% (30) as mild, providing actionable predictions for downstream validation. Users can further assess prediction confidence by examining the raw mutational sensitivity score relative to the protein-wide distribution (**Fig. 2A**).

### Decrypting intrinsically disordered regions

Predicting variant effects in disordered regions remains challenging. While ProteoCast achieved comparable overall accuracy on disordered (pLDDT<70) and ordered ClinVar mutations (0.84 vs 0.81), sensitivity for pathogenic variants drops substantially in disordered regions (Δrecall = 0.19), as expected given their weaker evolutionary constraints. ProteoCast segmentation approach addresses this challenge by identifying functional elements within disordered regions that evolve under locally stronger constraints than their background. We illustrate below how this enables detection of pathogenic variants and functional motifs that would otherwise be missed.

In the TERF1-interacting nuclear factor 2 TINF2, ProteoCast identified the known binding sites for TRF1 and HP1*γ* as more sensitive to mutations than their surroundings (**Fig. 2B**, purple segments). Furthermore, the known pathogenic mutations falling within the unstructured part of the protein are concentrated in HP1*γ*’s binding site (**Fig. 2B**, plain dots), while benign mutations are located in segments with lower sensitivity (**Fig. 2B**, circles). In the cyclin-dependent kinase inhibitor p27^*Kip*1^, ProteoCast identified four segments with elevated sensitivity, all corresponding to experimentally validated functional motifs (**Fig. 2C**): two degrons (residues 1-3 and 183-190), an AKT1 binding site (152-160), and a nuclear localisation signal (NLS, 153-166). More specifically, ProteoCast precisely highlighted the four critical basic residues (K153, R154, K165, R166) essential for nuclear import, including R166 whose mutation to alanine completely abolishes nuclear import^45^. By contrast, the known 14-3-3 binding site (residues 196-198) showed low mutational sensitivity and was not flagged by ProteoCast. This aligns with the peptide’s low binding affinity^46^ and suggests limited physiological relevance – a hypothesis supported by ProteoCast’s successful detection of the high-affinity 14-3-3 motif in KCNK3 (residues 391-394).

Beyond these case studies, ProteoCast shows enrichment for functional sites across proteomes. Specifically, 186 phosphorylation sites, localisation signals, degradation signals, SUMO sites, and interaction motifs in yeast, out of a total of ∼ 500, fall within segments identified by ProteoCast as having elevated mutational sensitivity. This represents twice the number detected by phylo-HMM over a dozen closely related species^17^. Together, these results demonstrate that ProteoCast can help users identify functionally constrained sites and discriminate between pathogenic and benign variants in disordered protein regions.

### Tuning the predictions to 3D conformations

ProteoCast leverages 3D structural data to tune predictions to specific functional states and to aid mechanistic interpretation of variant effects. Specifically, it uses per-residue relative solvent accessibility (RSA) values to weight the predicted evolutionary scores. As a result, residues buried in the input 3D structure are prioritised over exposed residues. This RSA-based modulation of the predictions yields an average Spearman rank correlation coefficient gain of Δ*ρ* = 0.0427 on the ProteinGym benchmark when using AlphaFold models (**Fig. 3A**). The RSA mostly contributes to improving prediction of thermodynamic stability, with Δ*ρ* > 0 in 97% of the assays (64 out of 66, **Fig. 3A**, red dots). Beyond stability predictions, solvent accessibility may improve predictions for binding assays when inputting complex structures, by putting more emphasis on the interacting surfaces. The SARS-CoV-2 spike glycoprotein provides an example where tuning the predictions to its conformation complexed with human ACE2 results in the interface residues gaining up to 89 ranks in predicted mutational sensitivity (**Fig. 3B**). These complex-informed predictions achieve high correlation with experimental measurements of the spike receptor binding domain variants’ affinities for ACE2^47^, with *ρ* = 0.62, compared to 0.54 when using the AlphaFold model and 0.30 when relying only on evolutionary scores. This case illustrates how simple structural descriptors such as solvent accessibility can enhance predictions when evolutionary signals are weak or ambiguous. When complexed conformations are not available, comparing the purely evolutionary-based predictions with those modulated by RSA on the monomeric state can help users identify residues important for protein interactions and functions, but not for stability. For instance, when inputting the monomeric structure of the human lipid-sensing transcription factor PPAR*γ*, two thirds of the 30 residues displaying the highest gain in mutational sensitivity upon excluding RSA establish molecular contacts with DNA and the substrate NCOA2 peptide (**Fig. 3C-D**, *Exposed*). By contrast, 26 out of the 30 most sensitive residues based on the RSA-modulated evolutionary scores either coordinate the zinc ions within the core of the zinc finger domains or establish intra-chain contacts (**Fig. 3C-D**, *Buried*).

**Figure 3.**
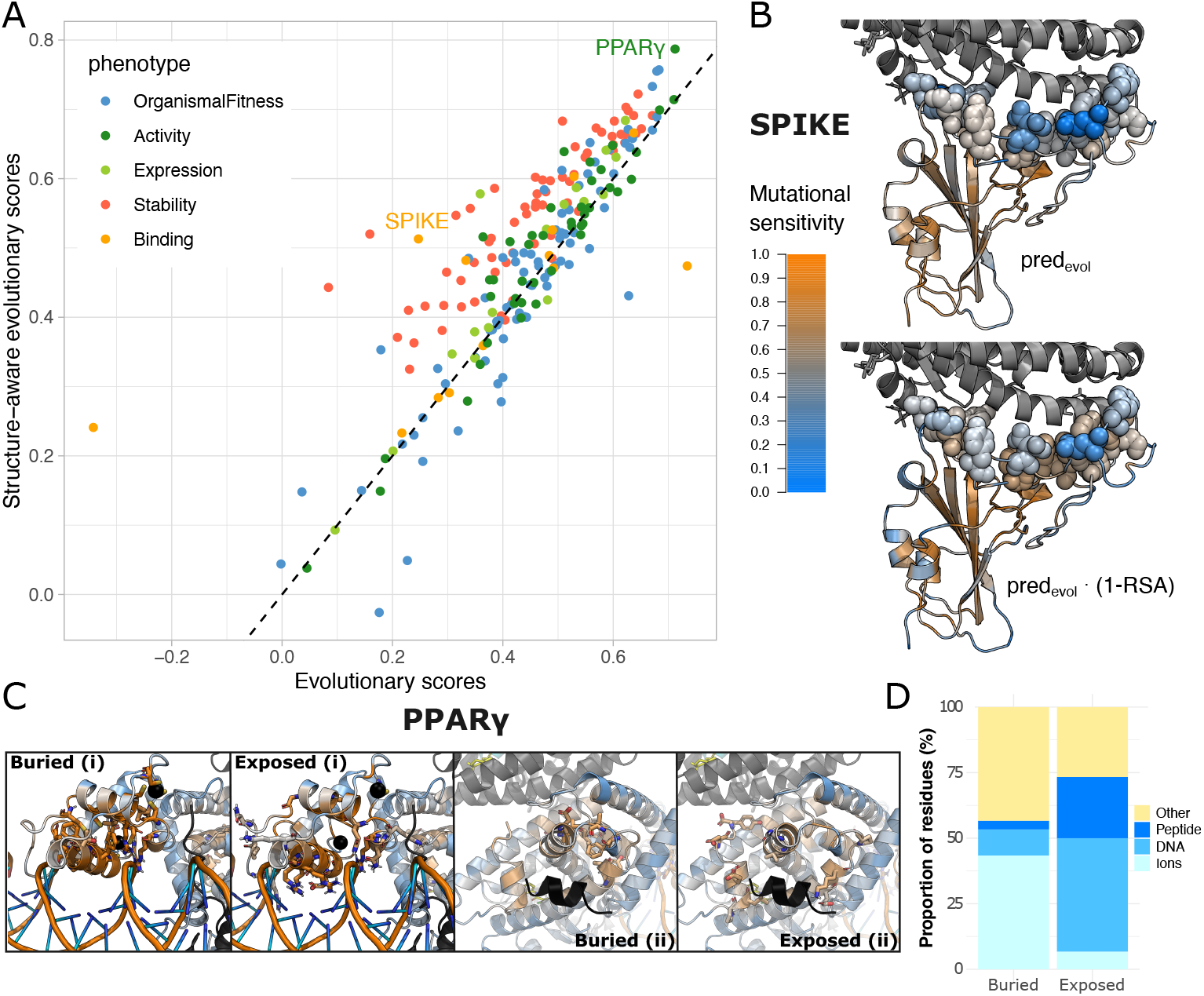
Structure-aware predictions. **A.** Comparison of Spearman rank correlation coefficients obtained computed on the 217 single-point mutational assays from the Prote-inGym benchmark when relying solely on evolutionary information (x-axis) or combining it with solvent accessibility (y-axis). **B-C**. Predictions mapped on 3D structures for the human lipid-sensing transcription factor PPAR*γ* (B, AlphaFold model superimposed with PDB complex 3E00^48^) and the spike protein from SARS-COV2 (C, PDB id: 6M0J^49^). The structures are displayed in cartoon with the residues of the query protein coloured according to their predicted mutational sensitivity. **B**. SPIKE residues at the interface with ACE2 (in grey) are highlighted in spheres. **C**. Close-up views of the DNA-binding (i) and ligand-binding (ii) domains of PPAR*γ*. Residues highlighted as sticks correspond to, Buried: the 30 residues with highest structure-aware sensitivity scores; Exposed: the 30 residues with highest evolutionary sensitivity and maximal sensitivity gain when excluding RSA. The zinc ions and the substrate NCOA2 peptide are in black while the DNA has its backbone in orange. **D**. Distribution of molecular contacts (<5^°^A) for the 30 top-ranked sensitive residues in PPAR*γ* from each category (Buried and Exposed, defined in panel C). Residues are classified by primary interaction partner: zinc ions, DNA, substrate NCOA2 peptide, or none of these.

## Conclusion

ProteoCast bridges the gap between cutting-edge variant effect prediction and practical accessibility for the research community. By combining transparent evolutionary analysis, structural awareness, and comprehensive quality controls within a user-friendly interface, it empowers researchers to interpret genetic variants and discover functional sites in both structured and disordered protein regions. Beyond ease of use, a key advantage of ProteoCast is its broader applicability: it covers virtually all variants, compared to methods like EVE that exclude a substantial fraction to achieve marginally higher sensitivity. Additionally, its segmentation strategy enables discrimination of constrained sites even within globally unconstrained protein contexts where evolutionary information is sparse. As genomic and proteomic data continue to expand, tools like ProteoCast will be essential for translating sequence variation into biological and clinical insights.

## Acknowledgments

This work was granted access to the HPC resources of the Institut Jacques Monod (IJM). It has been funded by ADAGIO – Ageing and Natural DeAth GenetIc cOntrollers (ANR-20-CE44-0010) and by the European Union (ERC, PROMISE, 101087830). Views and opinions expressed are however those of the author(s) only and do not necessarily reflect those of the European Union or the European Research Council. Neither the European Union nor the granting authority can be held responsible for them. We are grateful to Joël Marchand and Xavier Jeannin for their invaluable help in setting up the web server on the IJM cluster. Thanks to all contributing to open-source programming libraries, to all who deposit experimental data in public databases, to those who maintain these databases, and those who make methods available enriching experimental data.

## Author Contributions

Marina Abakarova: Methodology, Software, Validation, Investigation, Writing - Original Draft, Writing - Review & Editing, Visualization; Maria Ines Freiberger: Software, Validation, Investigation, Writing - Review & Editing, Visualization; Arnaud Liehrmann: Methodology, Validation, Writing - Review & Editing, Visualization; Michael Rera: Conceptualization, Methodology, Supervision, Funding acquisition, Writing - Review & Editing; Elodie Laine: Conceptualization, Methodology, Software, Validation, Investigation, Writing - Original Draft, Writing - Review & Editing, Supervision, Funding acquisition.

## References

[1] Pascal Notin, Aaron Kollasch, Daniel Ritter, Lood van Niekerk, Steffanie Paul, Han Spinner, Nathan Rollins, Ada Shaw, Rose Orenbuch, Ruben Weitzman, Jonathan Frazer, Mafalda Dias, Dinko Franceschi, Yarin Gal, and Debora Marks. ProteinGym: Large-scale benchmarks for protein fitness prediction and design. 36:64331–64379.

[2] Matsvei Tsishyn, Pauline Hermans, Fabrizio Pucci, and Marianne Rooman. Residue conservation and solvent accessibility are (almost) all you need for predicting mutational effects in proteins. Pages: 2025.02.03.636212 Section: New Results.

[3] Pauline Hermans, Matsvei Tsishyn, Martin Schwersensky, Marianne Rooman, and Fabrizio Pucci. Exploring evolution to enhance mutational stability prediction. Pages: 2024.05.28.596203 Section: New Results.

[4] Elodie Laine, Yasaman Karami, and Alessandra Carbone. GEMME: A simple and fast global epistatic model predicting mutational effects. 36(11):2604–2619.

[5] Daniel Hesslow, Niccoló Zanichelli Pascal Notin, Iacopo Poli, and Debora Marks. Rita: a study on scaling up generative protein sequence models. arXiv preprint 2205.05789, 2022.

[6] Céline Marquet, Julius Schlensok, Marina Abakarova, Burkhard Rost, and Elodie Laine. Expert-guided protein language models enable accurate and blazingly fast fitness prediction. 40(11):btae621.

[7] Konstantina Tzavella, Adrian Diaz, Catharina Olsen, and Wim Vranken. Combining evolution and protein language models for an interpretable cancer driver mutation prediction with d2deep. Briefings in Bioinformatics, 26(1), 2024.

[8] Lukas Gerasimavicius, Sarah A. Teichmann, and Joseph A. Marsh. Leveraging protein structural information to improve variant effect prediction. 92:103023.

[9] Yang Tan, Ruilin Wang, Banghao Wu, Liang Hong, and Bingxin Zhou. Retrieval-enhanced mutation mastery: Augmenting zero-shot prediction of protein language model.

[10] Alex S Holehouse and Birthe B Kragelund. The molecular basis for cellular function of intrinsically disordered protein regions. Nature Reviews Molecular Cell Biology, 25(3):187–211, 2024.

[11] Peter Tompa, Norman E Davey, Toby J Gibson, and M Madan Babu. A million peptide motifs for the molecular biologist. Molecular cell, 55(2):161–169, 2014.

[12] Mohamed Fawzy and Joseph A Marsh. Assessing variant effect predictors and disease mechanisms in intrinsically disordered proteins. PLOS Computational Biology, 21(8):e1013400, 2025.

[13] Maryam Mehdiabadi, Andrea Del Conte, Maria Vittoria Nugnes, Maria Chiara Aspromonte, Silvio C.E. Tosatto, and Damiano Piovesan. Critical assessment of protein intrinsic disorder (caid) round 3 – predicting disorder in the era of protein language models.

[14] Peter Tompa. Intrinsically unstructured proteins evolve by repeat expansion. Bioessays, 25(9):847–855, 2003.

[15] Celeste J Brown, Sachiko Takayama, Andrew M Campen, Pam Vise, Thomas W Marshall, Christopher J Oldfield, Christopher J Williams, and A Keith Dunker. Evolutionary rate heterogeneity in proteins with long disordered regions. Journal of molecular evolution, 55(1), 2002.

[16] Norman E Davey, Denis C Shields, and Richard J Edwards. Masking residues using context-specific evolutionary conservation significantly improves short linear motif discovery. Bioinformatics, 25(4):443–450, 2009.

[17] Alex N. Nguyen Ba, Brian J. Yeh, Dewald van Dyk, Alan R. Davidson, Brenda J. Andrews, Eric L. Weiss, and Alan M. Moses. Proteome-wide discovery of evolutionary conserved sequences in disordered regions. 5(215):rs1–rs1. Publisher: American Association for the Advancement of Science.

[18] Norman E. Davey, Joanne L. Cowan, Denis C. Shields, Toby J. Gibson, Mark J. Coldwell, and Richard J. Edwards. SLiMPrints: conservation-based discovery of functional motif fingerprints in intrinsically disordered protein regions. 40(21):10628– 10641.

[19] Mihaly Varadi, Damian Bertoni, Paulyna Magana, Urmila Paramval, Ivanna Pidruchna, Malarvizhi Radhakrishnan, Maxim Tsenkov, Sreenath Nair, Milot Mirdita, Jingi Yeo, Oleg Kovalevskiy, Kathryn Tunyasuvunakool, Agata Laydon, Augustin Žídek, Hamish Tomlinson, Dhavanthi Hariharan, Josh Abrahamson, Tim Green, John Jumper, Ewan Birney, Martin Steinegger, Demis Hassabis, and Sameer Velankar. AlphaFold protein structure database in 2024: providing structure coverage for over 214 million protein sequences. 52:D368–D375.

[20] Helen M. Berman, John Westbrook, Zukang Feng, Gary Gilliland, T. N. Bhat, Helge Weissig, Ilya N. Shindyalov, and Philip E. Bourne. The protein data bank. 28(1):235– 242.

[21] Arnaud Liehrmann, Etienne Delannoy, Alexandra Launay-Avon, Elodie Gilbault, Olivier Loudet, Benoît Castandet, and Guillem Rigaill. DiffSegR: an RNA-seq data driven method for differential expression analysis using changepoint detection. 5(4):qad098.

[22] Arnaud Liehrmann, Guillem Rigaill, and Toby Dylan Hocking. Increased peak detection accuracy in over-dispersed ChIP-seq data with supervised segmentation models. 22(1):323.

[23] Stefan Engelen, Ladislas A. Trojan, Sophie Sacquin-Mora, Richard Lavery, and Alessandra Carbone. Joint evolutionary trees: A large-scale method to predict protein interfaces based on sequence sampling. 5(1):e1000267. Publisher: Public Library of Science.

[24] Olivier Lichtarge, Henry R. Bourne, and Fred E. Cohen. An evolutionary trace method defines binding surfaces common to protein families. 257(2):342–358.

[25] Marina Abakarova, Maria Ines Freiberger, Arnaud Liehrmann, Michael Rera, and Elodie Laine. Proteome-wide prediction of the functional impact of missense variants with proteocast. bioRxiv, pages 2025–02, 2025.

[26] Robert Maidstone, Toby Hocking, Guillem Rigaill, and Paul Fearnhead. On optimal multiple changepoint algorithms for large data. 27(2):519–533.

[27] Guillem Rigaill. fpopw: Weighted Segmentation using Functional Pruning and Optimal Partioning.

[28] Peter Hall, JW Kay, and DM Titterington. Asymptotically optimal difference-based estimation of variance in nonparametric regression. 77(3):521–528.

[29] John Jumper, Richard Evans, Alexander Pritzel, Tim Green, Michael Figurnov, Olaf Ronneberger, Kathryn Tunyasuvunakool, Russ Bates, Augustin Žídek, Anna Potapenko, Alex Bridgland, Clemens Meyer, Simon A. A. Kohl, Andrew J. Ballard, Andrew Cowie, Bernardino Romera-Paredes, Stanislav Nikolov, Rishub Jain, Jonas Adler, Trevor Back, Stig Petersen, David Reiman, Ellen Clancy, Michal Zielinski, Martin Steinegger, Michalina Pacholska, Tamas Berghammer, Sebastian Bodenstein, David Silver, Oriol Vinyals, Andrew W. Senior, Koray Kavukcuoglu, Pushmeet Kohli, and Demis Hassabis. Highly accurate protein structure prediction with AlphaFold. 596(7873):583–589. Publisher: Nature Publishing Group.

[30] Valerio Mariani, Marco Biasini, Alessandro Barbato, and Torsten Schwede. lddt: a local superposition-free score for comparing protein structures and models using distance difference tests. Bioinformatics, 29(21):2722–2728, 2013.

[31] Peter JA Cock, Tiago Antao, Jeffrey T Chang, Brad A Chapman, Cymon J Cox, Andrew Dalke, Iddo Friedberg, Thomas Hamelryck, Frank Kauff, Bartek Wilczynski, et al. Biopython: freely available python tools for computational molecular biology and bioinformatics. Bioinformatics, 25(11):1422, 2009.

[32] Marcin Wojdyr. GEMMI: A library for structural biology. 7(73):4200.

[33] Andrew Shrake and John A Rupley. Environment and exposure to solvent of protein atoms. lysozyme and insulin. Journal of molecular biology, 79(2):351–371, 1973.

[34] Melissa J. Landrum, Jennifer M. Lee, Mark Benson, Garth Brown, Chen Chao, Shanmuga Chitipiralla, Baoshan Gu, Jennifer Hart, Douglas Hoffman, Jeffrey Hoover, Wonhee Jang, Kenneth Katz, Michael Ovetsky, George Riley, Amanjeev Sethi, Ray Tully, Ricardo Villamarin-Salomon, Wendy Rubinstein, and Donna R. Maglott. ClinVar: public archive of interpretations of clinically relevant variants. 44:D862–D868.

[35] Arzu Öztürk Çolak, Steven J Marygold, Giulia Antonazzo, Helen Attrill, Damien Goutte-Gattat, Victoria K Jenkins, Beverley B Matthews, Gillian Millburn, Gilberto dos Santos, Christopher J Tabone, and FlyBase Consortium. FlyBase: updates to the drosophila genes and genomes database. 227(1):iyad211.

[36] Trudy F. C. Mackay, Stephen Richards, Eric A. Stone, Antonio Barbadilla, Julien F. Ayroles, Dianhui Zhu, Sònia Casillas, Yi Han, Michael M. Magwire, Julie M. Cridland, Mark F. Richardson, Robert R. H. Anholt, Maite Barrón, Crystal Bess, Kerstin Petra Blankenburg, Mary Anna Carbone, David Castellano, Lesley Chaboub, Laura Duncan, Zeke Harris, Mehwish Javaid, Joy Christina Jayaseelan, Shalini N. Jhangiani, Katherine W. Jordan, Fremiet Lara, Faye Lawrence, Sandra L. Lee, Pablo Librado, Raquel S. Linheiro, Richard F. Lyman, Aaron J. Mackey, Mala Munidasa, Donna Marie Muzny, Lynne Nazareth, Irene Newsham, Lora Perales, Ling-Ling Pu, Carson Qu, Miquel Rámia, Jeffrey G. Reid, Stephanie M. Rollmann, Julio Rozas, Nehad Saada, Lavanya Turlapati, Kim C. Worley, Yuan-Qing Wu, Akihiko Yamamoto, Yiming Zhu, Casey M. Bergman, Kevin R. Thornton, David Mittelman, and Richard A. Gibbs. The drosophila melanogaster genetic reference panel. 482(7384):173–178. Publisher: Nature Publishing Group.

[37] Joaquin C B Nunez, Marta Coronado-Zamora, Mathieu Gautier, Martin Kapun, Sonja Steindl, Lino Ometto, Katja Hoedjes, Julia Beets, R Axel W Wiberg, Giovanni R Mazzeo, David J Bass, Denys Radionov, Iryna Kozeretska, Mariia Zinchenko, Oleksandra Protsenko, Svitlana V Serga, Cristina Amor-Jimenez, Sònia Casillas, Alejandro Sánchez-Gracia, Aleksandra Patenkovic, Amanda Glaser-Schmitt, Antonio Barbadilla, Antonio J Buendia-Ruíz, Astra Clelia Bertelli, Balázs Kiss, Banu Sebnem Önder, Bélen Roldán Matrín, Bregje Wertheim, Candice Deschamps, Carlos E Arboleda-Bustos, Carlos Tinedo, Christian Feller, Christian Schlötterer, Clancy Lawler, Claudia Fricke, Cristina P Vieira, Cristina Vieira, Darren J Obbard, Dorcas Juana Orengo, Doris Vela, Eduardo Amat, Elgion Loreto, Envel Kerdaffrec, Esra Durmaz Mitchell, Eva Puerma, Fabian Staubach, M Florencia Camus, Hervé Colinet Jan Hrcek, Jesper Givskov Sørensen, Jessica Abbott, Joan Torro, John Parsch, Jorge Vieira, Jose Luis Olmo, Khalid Khfif, Krzysztof Wojciechowski, Lilian Madi-Ravazzi, Maaria Kankare, Mads F Schou, Emmanuel D Ladoukakis, M Josefa Gómez-Julián, M Luisa Espinosa-Jimenez, Maria Pilar Garcia Guerreiro, Maria-Eleni Parakatselaki, Marija Savic Veselinovic, Marija Tanaskovic, Marina Stamenkovic-Radak, Margot Paris, Marta Pascual, Michael G Ritchie, Michel Rera, Mihailo Jelić, Mina Hojat Ansari, Mina Rakic, Miriam Merenciano, Natalia Hernandes, Nazar Gora, Nicolas Rode, Omar Rota-Stabelli, Paloma Sepulveda, Patricia Gibert, Pau Carazo, Pinar Kohlmeier, Priscilla A Erickson, Renaud Vitalis, Jorge Roberto Torres, Sara Guirao-Rico, Sebastian E Ramos-Onsins, Silvana Castillo, Tânia F Paulo, Venera Tyukmaeva, Zahara Alonso, Vladimir E Alatortsev, Elena Pasyukova, Dmitry V Mukha, Dmitri A Petrov, Paul Schmidt, Thomas Flatt, Alan O Bergland, and Josefa Gonzalez. Footprints of worldwide adaptation in structured populations of drosophila melanogaster through the expanded DEST 2.0 genomic resource. 42(8):msaf132.

[38] Marina Abakarova, Céline Marquet, Michael Rera, Burkhard Rost, and Elodie Laine. Alignment-based protein mutational landscape prediction: Doing more with less. 15(11):evad201.

[39] Milot Mirdita, Konstantin Schütze, Yoshitaka Moriwaki, Lim Heo, Sergey Ovchinnikov, and Martin Steinegger. ColabFold: making protein folding accessible to all. 19(6):679–682.

[40] Timothy Truong Jr and Tristan Bepler. PoET: A generative model of protein families as sequences-of-sequences. 36:77379–77415.

[41] Pascal Notin, Lood Van Niekerk, Aaron W. Kollasch, Daniel Ritter, Yarin Gal, and Debora S. Marks. TranceptEVE: Combining family-specific and family-agnostic models of protein sequences for improved fitness prediction. Pages: 2022.12.07.519495 Section: New Results.

[42] Jonathan Frazer, Pascal Notin, Mafalda Dias, Aidan Gomez, Joseph K. Min, Kelly Brock, Yarin Gal, and Debora S. Marks. Disease variant prediction with deep generative models of evolutionary data. 599(7883):91–95. Number: 7883 Publisher: Nature Publishing Group.

[43] Manjeet Kumar, Sushama Michael, Jesús Alvarado-Valverde, András Zeke, Tamas Lazar, Juliana Glavina, Eszter Nagy-Kanta, Juan Mac Donagh, Zsofia E Kalman, Stefano Pascarelli, Nicolas Palopoli, László Dobson Carmen Florencia Suarez, Kim Van Roey, Izabella Krystkowiak, Juan Esteban Griffin, Anurag Nagpal, Rajesh Bhardwaj, Francesca Diella, Bálint Mészáros, Kellie Dean, Norman E Davey, Rita Pancsa, Lucía B Chemes, and Toby J Gibson. ELM—the eukaryotic linear motif resource—2024 update. 52:D442–D455.

[44] Silvia Canudas, Benjamin R Houghtaling, Monica Bhanot, Ghadir Sasa, Sharon A Savage, Alison A Bertuch, and Susan Smith. A role for heterochromatin protein 1γ at human telomeres. Genes & development, 25(17):1807–1819, 2011.

[45] Ying Zeng, Katsuya Hirano, Mayumi Hirano, Junji Nishimura, and Hideo Kanaide. Minimal requirements for the nuclear localization of p27kip1, a cyclin-dependent kinase inhibitor. Biochemical and biophysical research communications, 274(1):37– 42, 2000.

[46] Alessandro Paiardini, Patrizia Aducci, Laura Cervoni, Francesca Cutruzzolá, Cristina Di Lucente, Giacomo Janson, Stefano Pascarella, Serena Rinaldo, Sabina Visconti, and Lorenzo Camoni. The phytotoxin fusicoccin differently regulates 14-3-3 proteins association to mode iii targets. IUBMB life, 66(1):52–62, 2014.

[47] Tyler N Starr, Allison J Greaney, Sarah K Hilton, Daniel Ellis, Katharine HD Crawford, Adam S Dingens, Mary Jane Navarro, John E Bowen, M Alejandra Tortorici, Alexandra C Walls, et al. Deep mutational scanning of sars-cov-2 receptor binding domain reveals constraints on folding and ace2 binding. cell, 182(5):1295–1310, 2020.

[48] Vikas Chandra, Pengxiang Huang, Yoshitomo Hamuro, Srilatha Raghuram, Yongjun Wang, Thomas P Burris, and Fraydoon Rastinejad. Structure of the intact ppar-γ– rxr-α nuclear receptor complex on dna. Nature, 456(7220):350–356, 2008.

[49] Jun Lan, Jiwan Ge, Jinfang Yu, Sisi Shan, Huan Zhou, Shilong Fan, Qi Zhang, Xuanling Shi, Qisheng Wang, Linqi Zhang, et al. Structure of the sars-cov-2 spike receptor-binding domain bound to the ace2 receptor. nature, 581(7807):215–220, 2020.

